# No evidence for an effect of chronic boat noise on the fitness of reared water fleas

**DOI:** 10.1101/2022.11.20.517267

**Authors:** Loïc Prosnier, Emilie Rojas, Vincent Médoc

## Abstract

Among the numerous questions about human impacts on ecosystems, there is a growing interest for acoustic pollution. First studies on underwater acoustic pollution focused, and showed effects, on vertebrates’ behaviours. Knowledge on the effects on invertebrates is more limited and there is a huge lack concerning zooplankton species, although widely used as bioindicators in chemical pollution. Consequently, it is critical to assess the impact of noise on zooplankton’s fitness (survival and fecundity). Here, isolated water fleas, *Daphnia magna*, were reared from birth to death in the presence or absence of motorboat noises. Effects on lifespan and clonal offspring production (e.g., clutch size, number of offspring produced along life) were assessed and chronic exposure to boat noise did not affect *Daphnia*’s fitness. The spectral and temporal features of the sounds could explain the results. This study highlights the importance of integrating noise pollution into ecotoxicological research to understand, and prevent, human impacts on communities.

## Introduction

Freshwater ecosystems are vulnerable to many types of anthropogenic pollution (e.g., chemicals, light, radioactivity, nanopollution, sounds) (see for instance Longcore & Rich, 2004; André et al., 2011; Song et al., 2020; Jan et al., 2022) but the most documented to date remain chemical pollutions (e.g., industrial effluents, urban waste, pesticides, drugs) (Truhaut, 1977; Villeneuve & Garcia-Reyero, 2011). Toxicological studies have documented, in a comprehensive and accurate way, the effects of different types of pollutants (e.g., ion, heavy metals, drugs), exposure durations (acute or chronic), intensities (e.g., median Lethal Dose LD_50_), and the interactions between them and with environmental parameters (temperature, acidity, humidity, etc.). Those results have contributed to the general knowledge allowing the evaluation of other types of pollution.

This study focuses on acoustic – or noise – pollution described as a pervasive and omnipresent pollutant found in all ecosystems (terrestrial, marine, and freshwater) (Shannon et al., 2016; Popper & Hawkins, 2019; Kunc & Schmidt, 2019), and which represents a growing research topic (Williams et al., 2015; Slabbekoorn, 2019). Specifically, this study focuses on the effect of boat traffic, an important source of noise likely to threaten aquatic systems (Rountree et al., 2020; Duarte et al., 2021). Studies on noise pollution have mainly focused on behaviour (Richardson et al., 1985; Duarte et al., 2021). Contrary to many ecotoxicological studies since Truhaut (1977), it remains a gap in understanding how noise pollution affects individual fitness, i.e. survival and fecundity (Francis & Barber, 2013; Read et al., 2014).

Although the effects of noise on large invertebrates, such as decapods or bivalves, have recently received substantial interest (see the reviews of Popper et al. (2001) and Solé et al. (2023)), research largely neglected zooplanktonic invertebrates (Hawkins et al., 2015; Prosnier, 2022; Vereide & Kühn, 2023), despite their ecological importance and general use as bioindicators in ecotoxicology (Parmar et al., 2016). Although zooplankton do not possess hearing structures, they present mechanoreceptors that allow them to detect particle motion, the other component of a sound with pressure (Gassie et al., 1993; Buskey et al., 2002). For instance, Gassie et al. (1993) found that marine copepods (*Acartia fossae*) can detect water vibrations, and Buskey et al. (2002) showed that vibrations can lead to acceleration in individuals of *Acartia spp*.. Marine zooplankton (e.g., copepods) exposed to acute airguns show a reduction of survival (McCauley et al., 2017; Fields et al., 2019; Vereide et al., 2023). Copepods also show reduced foraging rate during boat noise exposure (Kühn et al., 2023). *Chaoborus flavicans* larvae, an important predator of zooplankton, show more body rotations, interpreted as an anti-predatory behaviour, when exposed to boat noise for the first time (Rojas et al., 2021). These works highlight that noise could affect the fitness and behaviour of zooplankton species. However long-term exposure has not been investigated yet except *D. magna* exposed to chronic broadband noise, that showed a lower speed and surprisingly a higher fitness (Prosnier, Rojas, Valéro, et al., 2022). Temporal pattern is an important acoustic parameter of noise (Francis & Barber, 2013) with chronic exposure at the one hand and acute exposure at the other (Duarte et al., 2021). Chronic exposure means a continuous or intermittent, regular or random sound (e.g., turbine, boat noise) whereas acute exposure represents a punctual sound (e.g., airgun) (Nichols et al., 2015; McCauley et al., 2017). The spectral (i.e., sound level of each frequency) and temporal patterns of noise are known to affect the behaviour and physiology of organisms in different ways. For instance fish are more affected by random noise than by continuous or regular noise (Nichols et al., 2015). These results are interpreted as an ability for vertebrates to habituate to some predictable long-term noise exposition (Rojas et al., 2021). Consequently, this raises the question of whether some results with unrealistic noise, such as continuous broadband noise (Prosnier, Rojas, Valéro, et al., 2022), could be extrapolated to real situation where organisms are exposed to boat noise, thus exposed to intermittent random noise (i.e., unpredictable) with high spectral variability.

The aim of this study was to investigate the effect of chronic exposure to motorboat noise (intermittent and random noise with spectral variability) on the fitness of the water flea *Daphnia magna*, a common zooplankton species widely used in ecotoxicology (Ebert, 2022). Previous studies found no change in their mobility when exposed to acute noise (Sabet et al., 2015, 2019), whereas prior experiment with chronic exposure to broadband noise (i.e., a continuous noise) showed alterations in both fitness and behaviour (Prosnier, Rojas, Valéro, et al., 2022). If arthropods react like fish, then intermittent boat noises (i.e., unpredictable) should be more impactful than continuous broadband noise (i.e., predictable). Consequently, motorboat noise should affect the fitness of *D. magna*. Absence to negative effects are expected according to various chronic and acute exposure experiments on invertebrates (Solé et al., 2023), but note that a positive effect was found for *D. magna* chronically exposed to broadband noise (Prosnier, Rojas, Valéro, et al., 2022).

## Material and Methods

### Collection and maintenance of organisms

*Daphnia magna* had been purchased from Aqualiment (Grand Est, France) and stored in a 20-L rearing aquarium, filled with aged tap water (physiochemical composition is available on Zenodo repository (Prosnier, Rojas, & Médoc, 2022)), for one month. They were reared at 18°C under a 12:12 light:dark cycle. *Daphnia magna* were fed, every two days, with 0.05g of algae (i.e., 736 mJ) with a mix of 80% of *Arthrospira platensis* and 20% of *Aphanizomenon flos-aquae* (Algo’nergy® Spiruline + Klamath). Note that *A. platensis* is known to have a C:N ratio between 4 and 8 (Walach et al., 1987; Griffen & Drake, 2009).

### Fecundity and mortality

Reproductive success and survival were measured during an experiment similar as done in Prosnier, Rojas, Valéro, et al. (2022). Gravid *D. magna* were collected from the rearing aquarium and isolated in 50-mL jars containing Volvic® water. Newborns (< 24h) were transferred individually into 150-mL (5.6 × 8.6 cm) glass microcosm, closed with a 0.3-mm mesh tissue allowing water flows and noise transmission (Fig. 1).

**Figure 1.**
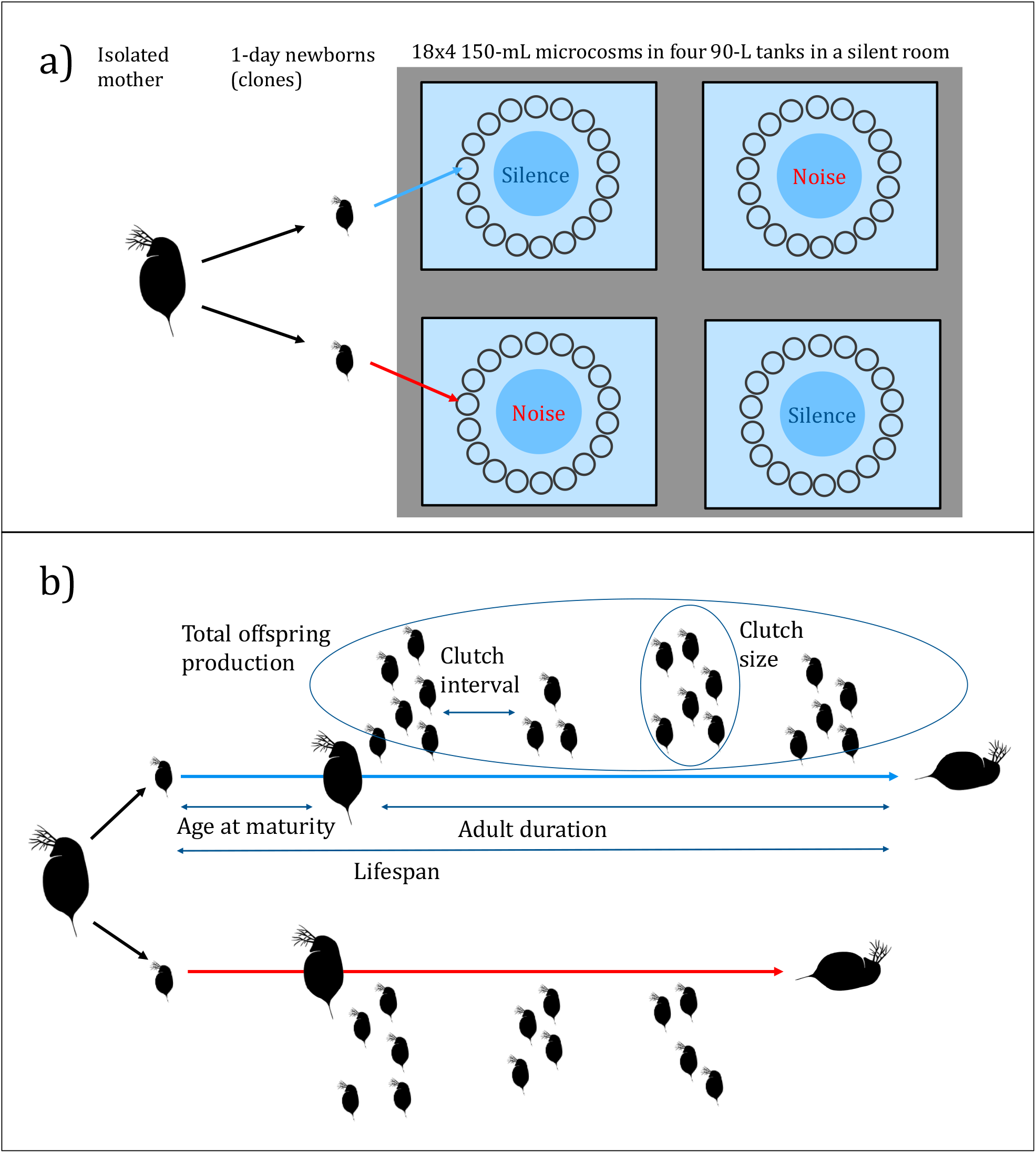
Setup. a) Experimental design. In the four tanks blue circles are loudspeakers, dark small circle are microcosms closed with net. b) Summary of all measures from birth to death of clonal individuals in the two treatments.

Eighteen glass microcosms were disposed at 20 cm of an underwater loudspeaker in four 150-L rectangular tanks (75 × 60 × 35 cm), filled with 90 L of aged tap water, at 20-22°C and under a 12:12 light:dark cycle. Silence was broadcasted in the two control tanks and a daily boat noise playlist (see below) used as treatment was broadcasted in the two other tanks. For each *D. magna* mother, half of the newborns were assigned to the control and the other half to the noise treatment, therefore individuals were clones in the two acoustic conditions. Each day, survival and newborns production were controlled, and when *D. magna* spawned, offspring were counted and removed. Every two days, individuals were fed with 2 mL of algae (1g/L) – note that both Griffen & Drake (2009) and Robinson et al. (2013) used a double quantity of *A. platensis*, but for populations starting with five to eighteen individuals –, and water was changed once a week. During the first eight days of the experiment (i.e., before the first hatching), dead *D. magna* were replaced by new newborns (isolated mothers were maintained in 50-mL jars during this initial period to be able to initiate new replicates with newborns) to increase the number of replicates. Experiments were performed with a total of 115 juveniles (58 control and 57 exposed to noise) coming from 25 mothers; almost half of the juveniles in each condition (26 in control and 23 in noise) reached maturity. The experiment lasted 46 days, from the birth of the first individual to the death of the last one (the oldest *D. magna* survived 39 days).

Based on daily survival and daily clutch, populational data were analysed using the Euler-Lotka equation (∑ *f*_*x*_*m*_*x*_*e*^−*rx*^ = 1), with *f*_*x*_ the fecundity at age *x, m*_*x*_ the survival at age *x*, and *r* the intrinsic rate of increase. This equation allows to calculate the reproductive output *R*_*0*_ (*R*_0_ = ∑ *f*_*x*_*m*_*x*_), the generation time 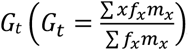, and the intrinsic rate of increase 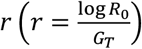 (Leung et al., 2007; Starke et al., 2021).

### Acoustic treatments

*Daphnia magna* were exposed to two acoustic treatments (see Rojas et al. (2021) for more details): a looped 1-h playlist without sound for the control (i.e., only the ambient noise), or boat noise for the treatment with a playlist including 15 recordings of motorboat sounds previously made in the Grangent lake (45°45′07.54″N, 4°25′56.47″E, Loire, France). Their intensity was modulated from 0 to -25 dB Re 1 μPa by 5 dB to create 75 sounds from 103 to 150 dB RMS Re 1 μPa – a naturally-occurring range of noise levels found in lakes (V. Médoc, pers. obs.) –, that were duplicated to obtain the total of 150 boat noises used in the experiment. The boat noise playlist was broadcasted from 9 a.m. to 6 p.m. (Fig. 2a). Both playlists (stereo WAV files) were generated using the Adobe Audition 2020 software (13.0.0.519, Adobe Systems Inc., Mountain View, CA, USA) and were played back using a Zoom® H4n recorder connected to an amplifier (DynaVox® CS-PA 1MK), and an underwater loudspeaker UW30 (Electro Voice®).

**Figure 2.**
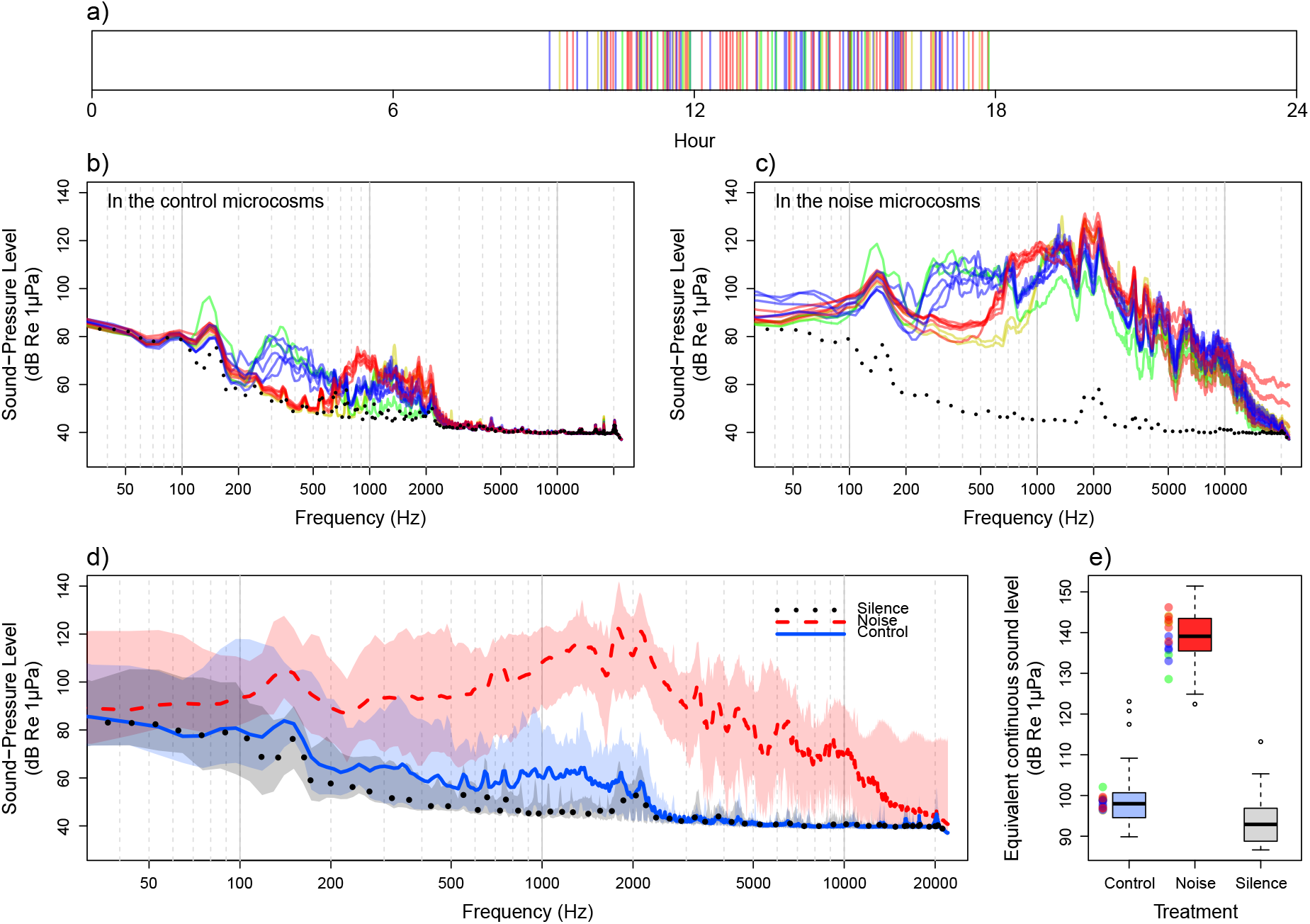
Acoustic treatments. a) 24-h temporal sequence of the broadcasted motorboat noises, from 9 a.m. to 6 p.m. Each vertical line represents a boat. b) Soundscapes of the control microcosms recorded in half microcosms with the records of the 15 boat sounds (solid lines), broadcasted at their maximal intensity in the noise tanks, and silence (dotted line). c) Soundscapes in the noise microcosms recorded in half of the microcosms during the broadcast of the15 boat sounds at their maximal intensity (solid lines) and silence (dotted line). d) Spectra in half of the microcosms. Thick lines are means for control (full blue line) and noise treatment (dashed red line) during the 15 boat broadcasts at their maximal intensity, and during the silence period (dotted black line). Shaded areas delimit the min and max Sound-Pressure Level. e) Sound levels in half of the microcosms. Central bars represent the median, boxes the interquartile range, and dots the outliers (> 1.5 times the interquartile range). Coloured dots are the sound levels in control and noise microcosms during the 15 boat broadcasts at their maximal intensity. The four colours (red, yellow, green, blue) correspond to four noise structure spectra that were visually determined (for instance red and yellow have low energy between 200 and 700 or 1000 Hz compared to green and blue boat noises).

To check spectra and noise levels in both control and noise microcosms, recordings were made (Fig. 2b-e) with a Zoom® H4n coupled to a hydrophone (Aquarian Audio H2A-HLR Hydrophone, frequency response from 10 Hz to 100 kHz) previously calibrated with a hydrophone (8104, Brüel & Kjær, Naerum, Denmark; sensitivity –205 dB re. V μPa− ; frequency response fro 0. Hz to 0 kHz) connected to a sound level meter (Bruël & Kjaer 2238 Mediator, Naerum, Denmark). Emitted boat noises were firstly corrected, using a one-third octave graphic equalizer (with Adobe Audition 2020), to make their spectrum closer to those of the original boat sound (Fig. 2). Boat noises were re-recorded only in half of the microcosms (in each tank) given that they were qualitatively and quantitatively similar due to the symmetry of the setup and after controlling with a broadband noise (Prosnier, Rojas, Valéro, et al., 2022). Note that the playback of boat noise was perceived in the control microcosms with intensities around 100 dB RMS Re 1 μPa, which was almost comparable to the sound level during the silence period (Fig. 2e) allowing to neglect noise transmission between the treatments. Particle motion cannot be measured due to the absence of adequate equipment, despite its importance for non-hearing species (Nedelec et al., 2016). However, Olivier et al. (2023) showed that results can still be qualitatively relevant when based solely on sound pressure level.

### Statistical analyses

Statistical analyses were performed using R (version 4.2.2) with a significant threshold at 5%. Data allowed to analyse separately the effects on mortality (death age and adult survival) and fecundity (age at maturity, clutch frequency, mean clutch size, and daily clutch size). The combination of mortality and fecundity was used as a proxy of fitness and quantified through total offspring production. Data were also studied at the population scale using the Euler-Lotka equation (but without statistical analysis due to absence of populational replicates). A survival analysis (Log-Rank test) was performed to compare survival (death age and adult duration) and age at maturity (first clutch age) between the control and noise treatments. For the fecundity parameters, only individuals that clutched at least once (i.e., that reached maturity) were considered in the analyses. Clutch frequency (i.e., mean time between two clutches) and mean clutch size were analysed using linear mixed-effects model, with tanks as random effect, thanks to the normal distribution of the data checked with a Shapiro test. The effect of both noise and age on daily clutch size was analysed by a type-II analysis of variance, completed with a pairwise Wilcoxon test between the treatments within each age. To test the effect of noise on the total number of clutches and offspring along life a generalized linear mixed-effects models was used, with tanks as random effect and a log function as the link function for Poisson distribution.

## Results

Chronic boat noise did not affect (Table A1) the survival of *D. magna* (p-value = 0.51, Fig. 3a) with a median survival of 4 days for the control and 5 days for the noise treatment. It did not affect the fecundity parameters, with clutch interval around 2.5 days (p-value = 0.24, Fig. 3b), mean clutch size around 10 offspring (p-value = 0.74, Fig. 3c) and age at maturity around 8 days (p-value = 0.65). Daily clutch size was not influenced by noise (noise: p-value = 0.89, noise x age: p-value = 0.35, pairwise: p-values > 0.38; Fig. A1), but changed with age (p-value = 0.003) with larger clutches at intermediate ages. However, taking into account the whole lifespan, there was an effect on the total number of clutches (p-value = 0.003) with 5 clutches for the control and 6 clutches for the noise treatment, and a tendency for higher total offspring production under boat noise exposure (p-value = 0.099, Fig. 3d,e), with on average 60 to 70 newborns.

**Figure 3.**
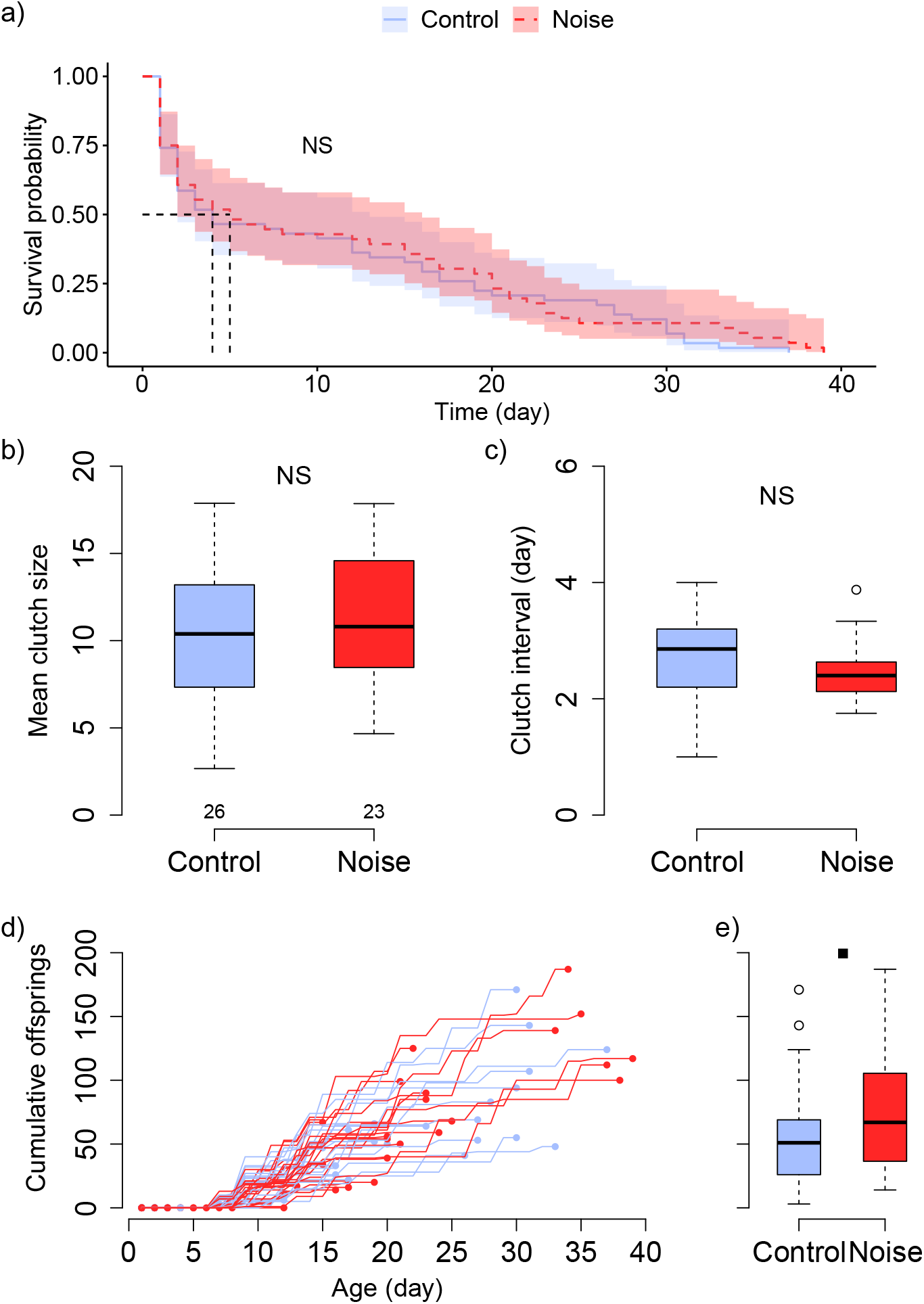
Effects of noise treatments on *Daphnia magna* survival and fecundity. a) Survival of *D. magna*; b) mean clutch size; c) clutch frequency; d) Cumulative offspring production along life; and e) total number of offspring during lifetime. Numbers in b) are the numbers of *D. magna* for the two treatments for b, c and e. a) Representation according to the Kaplan-Meier method; b-e) central bars represent the median, boxes the interquartile range, and dots the outliers (> 1.5 times the interquartile range). Statistical analysis: dot P < 0.1, * P < 0.05; ** P < 0.01; NS P>0.1. See Table A1 for statistical values.

The populational analysis done with the Lotka-Euler equation confirmed the tendancy on total offspring production with a reproductive output (*R*_*0*_) higher for the noise treatment with 63 offspring compared to the 54 offspring in control. Generation time (*G*_*T*_) was longer in the noise treatment, with 16.6 days, compared to the 15.2 days in the control. The combination of both led to an intrinsic rate of increase of 0.25 day^-1^ in the noise treatment compared to 0.26 day^-1^ in the control.

## Discussion

This study investigated the effect of exposure to chronic boat noise on the fitness of the water flea *Daphnia magna*. Contrary to what expected, no effect on the survival and fecundity parameters was observed, which differs from results obtained with exposure to another type of chronic noise (Prosnier, Rojas, Valéro, et al., 2022). Such difference in the results might be due to variations in the temporal and spectral features of the noise. Moreover, as one of the first experiments on chronic noise, it raises questions on the interactions with other pollutants, and on the effects on complex communities.

Chronic boat noise had no effect on *Daphnia magna*’s fecundity and survival. These results are opposite to those on acute and intense exposition of copepods to airgun. McCauley et al. (2017) observed high mortality for numerous marine zooplankton species including copepods following the passage of a boat equipped with an airgun. Fields et al. (2019) showed that airgun exposure leads to increased mortality in *Calanus finmarchicus* within an hour. Our results are also opposite to those of Prosnier, Rojas, Valéro, et al. (2022), where an exposition to a continuous broadband noise leads to a counter-intuitive increase in *D. magna*’s fitness, with higher survival and greater clutch size. Here, although boat noise had no statistical effect on the survival and fecundity parameters, there was a tendency for a higher total offspring production with noise, which would be consistent with the results of Prosnier, Rojas, Valéro, et al. (2022). However, at a population scale, the Euler-Lotka equation suggests a tendency for a lower growth when exposed to boat noise due to a shorter generational time. Such differences in the magnitude of the effects suggest that response of zooplankton to chronic noise pollution could depend on the temporal and spectral structure of noise. Indeed, in Prosnier, Rojas, Valéro, et al. (2022), the broadband noise broadcasted was continuous and at high level (130 dB RMS re 1 μPa) for all frequencies (from 0.1 to 20 kHz), whereas, in this study, sounds with a random temporal pattern (a total of 2h of noise *per* day) and different spectra (some boat noises with low intensity between 200 and 800 Hz) were used at various levels (from 108 to 136 dB RMS re 1 μPa). It is already known that, in hearing vertebrates (i.e., with dedicated organs to detect sound pressure variation as inner), animals respond differently to chronic noise pollution depending on temporal variations (continuous, regular, random), spectral variations (i.e., variation in the frequencies), and if noise is predictable or not (Francis & Barber, 2013). Nichols et al. (2015) showed that fish were more stressed (higher cortisol concentration) with a higher noise level (from 126 to 141 dB RMS re 1 μPa), and under intermittent random noise (i.e., unpredictable) compared to continuous noise. However, the review of de Jong et al. (2020) on noise effects on fecundity revealed that continuous noise with spectral variations, such as boat noise, was more prone to impact physiological markers (cortisol level, ventilation rates and metabolic rate) and behaviour (startle and freeze responses, horizontal and vertical avoidances). Another study on zebrafish larvae showed a negative effect of a continuous white noise on survival and a higher cortisol concentration (Lara & Vasconcelos, 2021). Thus, it seems that for hearing vertebrates, depending on the developmental stage and the considered characteristic (fitness, behavior, or physiology), both temporal and spectral characteristics need to be considered. For zooplankton, little is known about the importance of noise characteristics. Here, the comparison with Lara & Vasconcelos (2021) and Prosnier, Rojas, Valéro, et al. (2022) suggests that negative effects are higher with continuous noise, contrary what is reported with fishes.

Despite there was no effect on fecundity, it would be necessary to focus on the offspring coming from mothers exposed to noise. Here, there was no qualitative effect reported during the current experiments, i.e., all the water fleas produced seemed viable and mobile. More, there was no increase in the mortality of newborns due to noise, and no effect was reported on the size of *D. magna* exposed to chronic noise (Prosnier, Rojas, Valéro, et al., 2022). This would be consistent with the study of Day et al. (2016) where exposure to air gun did not affect the embryonic development of the spiny lobster *Jasus edwardsii* (Decapodae). However, airgun exposition reduces growth and development stage of *Acartia tonsa* nauplii (Vereide et al., 2023). This aspect seems important as it is known that stress on mother and early stages can affect daphnia’s develop ent (Mittmann et al., 2014; Mushegian et al., 2016) and that effects can differ across generations (Campos et al., 2016). Consequently, impact studies on noise should focus on embryonic development and perform multigenerational experiments to determine the long-term effects of chronic exposure resulting from embryonic misdevelopment (Mushegian et al., 2016), maternal effects (Radersma et al., 2018), and acclimatation or adaptation (Ringot et al., 2018; Abdullahi et al., 2022).

An interesting question is to consider the effect of noise as part of cocktails of pollutants. It is now a common question in ecotoxicology to ask whether stressors (e.g., chemical pollution, temperature, food quality) act synergistically (Altshuler et al., 2011). For instance, Starke et al. (2021) showed that food quality impacts more *D. pulicaria* at some higher temperature due to the increased metabolism. Prosnier et al. (2015) modelled the antagonistic effect of copper and nutrient enrichment on the *Daphnia* - algae interaction. Regarding noise, Stenton et al. (2022) showed both synergetic and antagonistic effects of noise and cadmium exposure on Norway lobster larvae (*Nephrops norvegicus*), depending of the considered parameters (survival, development, oxidative indicators). Another, McMahon et al. (2017) studied the interactive effects of light and noise pollutions on a frog-parasite interaction. They showed that light reduced frog-biting midge (*Corethrella* spp.) abundance at low noise level, whereas there was no midge at high noise level. It could be also useful to investigate whether all the unwanted noises produced by many experimental setups (to control light, temperature, oxygenation, and food) interact with the stressors studied and influence the results. For instance, in the present study there was a very high mortality in *D. magna* juveniles compared to similar studies (Parisot et al., 2015; Prosnier, Loeuille, et al., 2023). It suggests suboptimal conditions (i.e., other stressor than noise), such as a lack of food – for instance Serra et al. (2020) fed *D. magna* daily for the seven first day –, which may have affected the outcomes through the masking of effects for example. However, on the other side, suboptimal conditions could make individuals more prone to be affected by an additional stress like noise. Note that, with the same suboptimal conditions, Prosnier, Rojas, Valéro, et al. (2022) obtained a significant difference between the control and noise treatments. The recent *Larvosonic* system, developed by Olivier et al. (2023), could be useful to study the impact of noise on zooplankton with a better control of the environmental conditions.

This study is a first step in our understanding of the importance of noise patterns for invertebrates (i.e., predictable versus unpredictable noise) in comparison with vertebrates. To go further, we need more information about noise perception (i.e., mechanoreception and involved gene) and sensory integration, that could explain the mentioned differences between vertebrates and invertebrates and seem largely unexplored (Gassie et al., 1993; Popper et al., 2001). Understanding the various reactions of vertebrates and invertebrates in terms of behavior, but also in terms of fitness is mandatory to study how noise could affect complex communities (Francis et al., 2009; Slabbekoorn & Halfwerk, 2009; Slabbekoorn, 2019). For instance, in a freshwater community, unpredictable noise should affect more fishes, at top trophic levels, than invertebrates. But if there are various effects within zooplankton community, as expected, leading to bottom-up effects (Rojas, Desjonquères, et al., 2023; Rojas, Gouret, et al., 2023). Zooplankton is highly diverse and predatory species might react differently than their zooplanktonic prey. Moreover, in a community, pollutants can alter fitness directly (as in this study) but also indirectly through change in the vulnerability to natural enemies (Read et al., 2014). For instance, noise does not affect frog abundance but reduces that of their parasite (McMahon et al., 2017). The need for more research on invertebrates and fitness impacts, particularly in arthropods, is also true for terrestrial communities (Morley et al., 2014). Thus, a more general overview on the response of invertebrates to anthropogenic noises should be beneficial to mitigate the impacts (Francis & Barber, 2013).

## Acknowledgments

The authors would like to thank all the people who contributed to the success of this experiment: Nicolas Boyer and Aurélie Pradeau for *Daphnia* rearing and providing material, Léo Papet, Joël Attia, and Jérémy Rouch for the acoustic treatments and analysis and for providing material, Olivier Valéro for sound recording, Marilyn Beauchaud and Paolo Fonseca for acoustic calibration, Théophile Turco for help in the analyses, and the EYD (ENES Young Discussion) for the useful discussions. Authors also thanks Marie-Agnès Coutellec and the anonymous reviewer for their useful comments, and Claudia Cosio for its recommendation (Cosio, 2023).

Note that this article has been published as a chapter of *The effects of noise on aquatic life: Principles and Practical Considerations* (Popper et al., 2023). DOI: <u>10.1007/978-3-031-10417-6_129-2</u> (Prosnier, Rojas, et al., 2023)

## Data, script and code availability

Data, script and code are available on Zenodo. DOI: 10.5281/zenodo.7775919 (Prosnier, Rojas, & Médoc, 2022)

## Supplementary information

Supplementary information is available after the references:

- Appendix: Table of statistics and supplementary figure

## Appendix: Table of statistics and supplementary figure

**Table A1.** Statistical results of chronic boat noises effects on fecundity and mortality of *Daphnia magna* (Fig. 3)

|  | Mortality |  | Reproduction |  |  | Fitness |  |
| --- | --- | --- | --- | --- | --- | --- | --- |
|  | Survival | Adult duration | Age at maturity | Clutch interval | Mean clutch size | Number of clutches | Number of offspring |
| df | 1 | 1 | 1 | 1 | 1 | 1 | 1 |
| $\chi^2/W$ | 0.4 | 1.6 | 0.2 | 1.4029 | -0.54276 | 3.8197 | 1.8263 |
| P-value | 0.51 | 0.21 | 0.65 | 0.236 | 0.737 | **0.031** | 0.099 |
| R2 | 0.0041 | 0.03 | 0.0046 | 0.03 | 0.007 | 0.096 | 0.327 |

**Figure A1.**
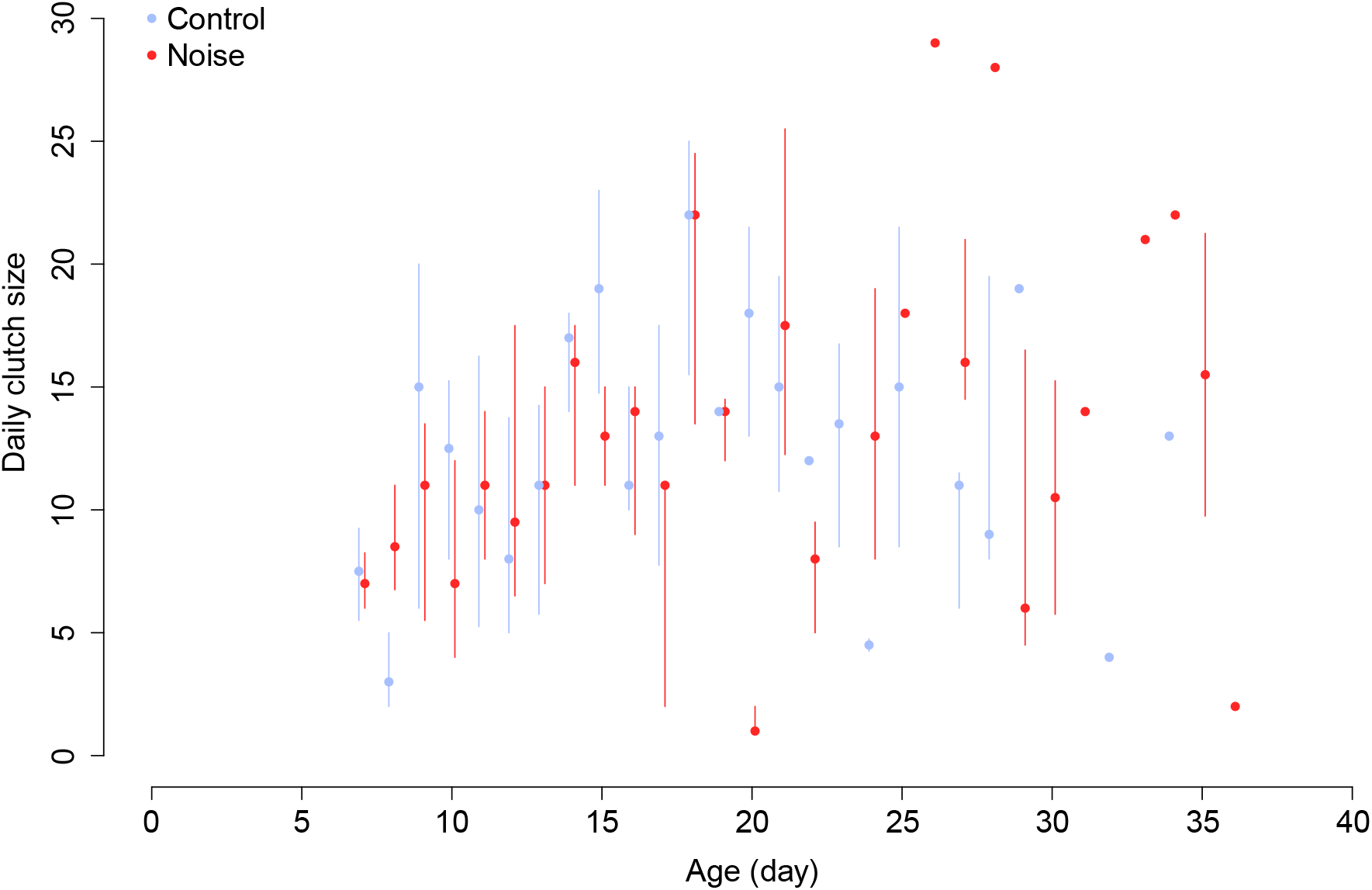
Effects of noise treatments on *Daphnia magna* daily clutch size. Dots represent the median of the daily clutch size and lines are interquartile ranges. See Fig. 3a,d as complement for the number of individuals at each age. Note that there could be only one clutch for an age (i.e., no interquartile lines) or clutches only for one treatment (i.e., only one point for an age).

